# Synaptic Synchronization-Based Learning of Pattern Separation in Self-Organizing Probabilistic Spiking Neural Networks

**DOI:** 10.1101/2025.11.08.687345

**Authors:** Faramarz Faghihi, Ahmed Moustafa, Samuel Neymotin

## Abstract

Neuroscience-inspired neural networks bridge biology and technology, offering powerful tools to model brain function while enabling adaptive, efficient control in robotics. In this work, we present a neuroscience-inspired synaptic learning rule based on the synchronization of synaptic inputs to single excitatory neurons within a feedforward spiking neural network. The model consists of three excitatory layers and two feedback inhibitory layers, with initially low connection probabilities and weak synaptic weights assigned to the excitatory neurons. Under an unsupervised learning paradigm, stimulus patterns were presented to the network, allowing synaptic weights and connectivity to evolve dynamically across training trials. We investigated how these dynamics depended on feedback inhibition intensity and identified conditions under which the network achieved stable activity. Furthermore, we evaluated the model’s pattern separation efficacy and its relationship to network dynamics. The results highlight the critical role of feedback inhibition in both stabilizing the network and enhancing pattern separation. In particular, results show balanced synchronization between excitatory and inhibitory populations maximizes separation efficacy. Beyond providing a novel computational framework for understanding information processing in neural systems, this model also offers insights into cognitive disorders associated with impaired inhibition and pattern separation, such as autism and schizophrenia. Finally, we embedded the trained network within a simulated agent navigating a two-dimensional environment, where it was tasked with identifying a trained stimulus as an obstacle and avoiding it. The model offers a framework for advancing cognitive robotics by enabling novel approaches that mimic natural intelligence and support the learning of complex environmental patterns.

## Introduction

Neuroscience-inspired artificial intelligence (AI) leverages the structure and function of the brain to design adaptive and efficient computational models. By mimicking neural processes such as spiking activity, synaptic plasticity, and network organization, researchers aim to build AI systems that learn and process information in brain-like ways. This synergy between neuroscience and AI has driven innovations such as Spiking Neural Networks (SNNs), neuromorphic hardware, and brain-inspired learning algorithms, offering pathways to energy-efficient computing and more human-like intelligence^**1**^.

SNNs model realistic neural activity with high temporal precision, and allow study of neural learning, plasticity, and network dynamics^**2,3**^ SNN’s event-driven nature makes them useful for neuromorphic hardware, allowing robots to perform tasks such as navigation, object recognition, and motor control in a way that mimics biological intelligence^**4, 5**^.

Probabilistic Spiking Neural Networks (PSNNs) integrate the principles of probabilistic computation with the dynamics of spiking neurons. Instead of producing deterministic outputs given an input, these networks model uncertainty by incorporating randomness in neural responses such as spike timing, firing probability, or synaptic transmission^**6,7**.^ This stochastic behavior enables PSNNs to represent and manipulate uncertainty just as biological neural systems do in tasks like perception, prediction, and decision-making^**8,9**^. PSNNs offer a computationally efficient alternative to traditional integrate-and-fire (I&F) models, particularly in large-scale or inference-driven applications. Unlike I&F neurons, which require continuous tracking of membrane potentials through numerical integration, probabilistic models generate spikes based on simplified statistical rules, often driven by input rates or instantaneous likelihoods. While I&F models provide greater biological realism and temporal precision, probabilistic SNNs are increasingly used in neuromorphic computing and probabilistic inference tasks where efficiency and scalability are key priorities^**10,11,12**^.

Balance between excitatory (E) and inhibitory (I) neuronal activity (E/I balance) is essential for proper brain function^**13,14**^. This balance ensures that neural circuits operate within an optimal range preventing excessive excitation that can lead to hyperactivity or seizures, and avoiding excessive inhibition that can suppress necessary neural signaling^**15,16**^. Disruptions to balanced synchronization between inhibitory and excitatory neurons can arise from altered synaptic strengths, dysregulation of neurotransmitters, and neurodevelopmental abnormalities affecting inhibitory interneuron development^**17,18**^. E/I imbalances, impair the fine-tuned neural dynamics required for efficient information processing^**19,20**^. Cognitive disorders linked to E/I imbalance manifest as deficits in decision-making, memory dysfunction, and impaired sensory processing. As an example, schizophrenia is associated with disrupted E/I balance leading to symptoms such as impaired working memory^**21**^. Studies have shown that disrupted E/I balance particularly in resting-state brain activity, contributing to memory loss and cognitive decline in patients with Alzheimer^**22**^. Therefore, understanding these mechanisms provides important insights for therapeutic strategies aiming to restore E/I balance and improve cognitive function in neurological and psychiatric disorders.

Pattern separation is a neural computation that enables the brain to distinguish between similar inputs or experiences by creating distinct representations^**23,24,25**^. This function, primarily associated with the hippocampus particularly the dentate gyrus is essential for episodic memory, spatial navigation, and learning^**26,27,28**^. It ensures that overlapping stimuli or events do not interfere with one another, thus preventing memory confusion and supporting accurate recall. Computational studies and simulations have been developed to study the parameters that are not accessible in the experimental approaches^**29,30,31**^. These parameters include biophysical features of the neurons, different synaptic learning rules, neuronal population size, connectivity of neurons, etc.^**32,33,34**^. In cognitive disorders such as Alzheimer’s disease and schizophrenia, impairments in pattern separation are commonly observed. These deficits can lead to difficulties in distinguishing between similar situations or memories, resulting in confusion, false recognition, problems with memory retrieval, or reduced learning capacity^**35,36**^. The balance between excitatory (E) and inhibitory (I) neuronal activity is crucial for efficient pattern separation in biological neural systems. In regions such as the hippocampus particularly the dentate gyrus lateral inhibition and well-tuned E/I ratios help to separate similar input patterns, reducing overlap and enhancing discrimination. When excitation and inhibition are out of balance either by weakening inhibition or by excessive excitation the capacity for separating similar inputs deteriorates, leading to less reliable memory representations and impaired cognitive performance^**30,37**^.

In artificial neural networks, non-Hebbian learning rules refer to mechanisms of synaptic weight adjustment that do not follow the classical Hebbian principle of “cells that fire together wire together”. Instead, these rules rely on alternative factors such as global error signals, activity from surrounding neurons, or temporal constraints^**38,39,40**^. A prominent example is the backpropagation algorithm, which updates weights based on the gradient of a loss function with respect to network output^**41,42**^. Unlike classic Hebbian rules that rely on the precise co-activation of pre- and postsynaptic neurons, these mechanisms are governed by global feedback signals or intrinsic activity constraints, enabling broader regulation of synaptic efficacy^**43,44**^. These rules allow artificial systems to learn more flexibly and robustly than would be possible with Hebbian mechanisms alone, supporting applications such as pattern recognition, motor control, and adaptive behavior in dynamic environments^**45,46,47**^.

Synchronization-based information transfer refers to the process by which neurons receive and interpret information not just from the rate of incoming spikes, but from their precise timing and temporal coordination^**48,49,50**^. When multiple presynaptic neurons fire in a synchronized manner, their combined inputs can summate more effectively at the postsynaptic neuron, making it more likely to generate a spike. This temporal alignment enhances the signal-to-noise ratio and allows single neurons to act as coincidence detectors, responding selectively to coordinated patterns of input^**51,52,53**^. This mechanism plays a critical role in various neural computations, including sensory processing, attentional modulation, and temporal coding. For example, in the visual and auditory systems, the synchronization of input spikes can help encode stimulus features with high temporal precision. In cortical circuits, gamma-band synchronization (30–80 Hz) is thought to dynamically route information across neural populations, allowing individual neurons to selectively respond to inputs from specific, coherently active groups^**50,54,55**^.

During the early stages of neurodevelopment, the formation of synaptic connections is largely guided by activity-independent processes such as molecular signaling, structural plasticity, and spontaneous neural activity, rather than by activity-dependent mechanisms like Hebbian learning. At this stage, Hebbian learning is not efficient because the sparse and immature connectivity does not yet provide sufficient coincident activity to drive synaptic strengthening. Instead, mechanisms based on synchronization and structural remodeling are likely to play a dominant role in establishing the initial network architecture^**56,57,58,59**^. Only after a sufficient level of connectivity has been achieved does Hebbian learning become more effective in refining and stabilizing synaptic connections.

Motivated by this biological perspective, we propose a synchronization-based learning rule designed to capture aspects of early developmental synaptic dynamics. In this work, we propose a new learning rule inspired by early stages of neurodevelopment specifically, the wiring of neural circuits in the growing brain of an infant. This rule is implemented in a probabilistic, three-layered feedback spiking neural network that also includes two inhibitory layers functioning as a probabilistic feedback mechanism. Drawing on biologically inspired wiring rules and the proposed synaptic learning mechanism, we present a novel neural network model capable of self-organizing its inter-layer connections in response to input patterns during the training phase. The network autonomously adapts and structures its connectivity without supervision.

We describe the architecture of the neural network and its underlying probabilistic mechanisms, followed by a series of unsupervised learning experiments focused on basic pattern separation. These experiments investigate the influence of various parameter values, particularly the intensity of inhibition. Finally, the optimized neural network is integrated into a simulated robot tasked with navigating a two-dimensional environment while avoiding specific stimuli it has been previously trained to recognize.

## Model and Methods

### Network architecture

Inspired by cortical architecture and hippocampal circuits, we developed a three-layered feedback Probabilistic Spiking Neural Network (PSNN) composed of both excitatory and inhibitory neurons (Fig. 1A). The first layer consists of 100 sensory neurons that encode stimulus.

**Fig. 1.**
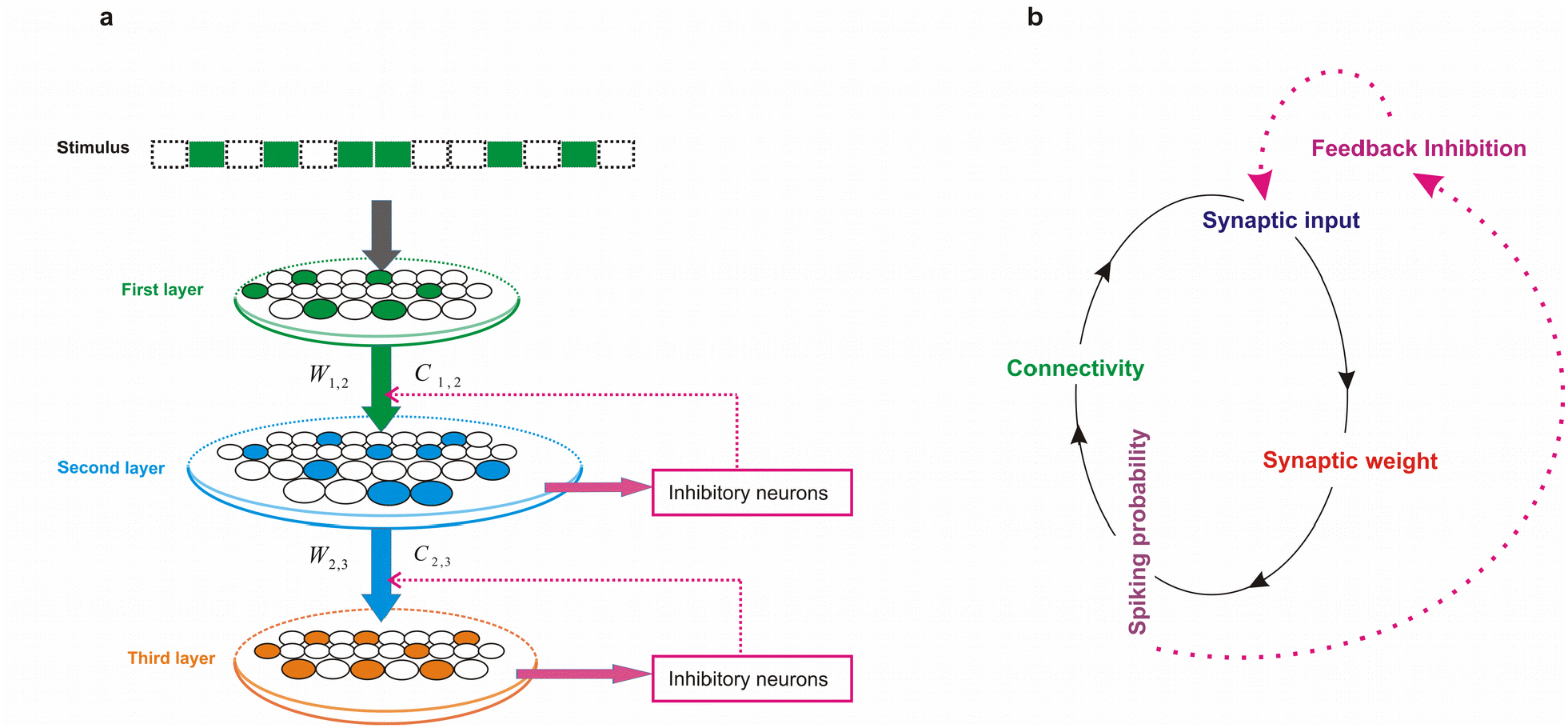
**a. Neural network architecture.** The model consists of one stimulus layer, three probabilistic excitatory spiking layers, and two inhibitory layers, each fully connected to its corresponding excitatory layer. Within this feedback architecture, every excitatory neuron is paired with a specific inhibitory neuron. Synaptic weights and connectivity values were initialized at 0.05 and dynamically updated during training. Neural activity in the final excitatory layer was used to evaluate pattern separation efficacy in simulations. **b. Schematic of neural dynamics during training**. At the end of each trial, inputs to a neuron modify its synaptic weights based on the synchronization level of incoming spikes. The neuron’s spiking probability is influenced by changes in excitation, which depend on both inputs and synaptic weights. Connectivity is updated according to the postsynaptic neuron’s spiking probability. Feedback inhibition further regulates synaptic input strength based on the inhibition intensity parameter.

In the experimental setup, stimulus pattern is defined as fixed subsets of neuron indices, each representing a distinct input condition. During training, these patterns are presented to the network as 100 time bins, with spike trains generated over a duration of 500 milliseconds in each time bin. Within each stimulus pattern, only the neurons assigned to the pattern are active; the remaining neurons remain quiescent. Spike generation for each active neuron follows a Bernoulli process with a firing probability p ∈ [0,1], independently sampled for each neuron at the beginning of the simulation and held constant across presentations. Thus, an active neuron emits a spike (denoted as 1) with probability p, and remains silent (0) otherwise. For the pattern separation experiments, all stimulus patterns are constructed to be fully disjoint, i.e., with no overlapping active neurons between any pair of patterns. This ensures orthogonality in the input space, enabling a rigorous assessment of the network’s capacity to form distinct internal representations for non-overlapping inputs.

The second and third layers are probabilistic spiking neurons whose activity is determined by the model dynamics. The network comprises 300 in the second, and 100 in the third. Both the second and third layers include inhibitory neurons, with each excitatory neuron paired with a corresponding inhibitory neuron in a one-to-one connection scheme. Inhibitory neurons do not inhibit each other; instead, each inhibitory neuron connects to a single excitatory neuron and provides probabilistic feedback inhibition through synaptic connections back to that excitatory neuron.

The synaptic weight between each excitatory neuron and its paired inhibitory neuron was fixed at 1 throughout the training and the simulations.

At the beginning of training, the probability of forming a connection between neurons in adjacent layers was set to 0.05. Similarly, initial synaptic weights were set to 0.05. Both connectivity and weights were dynamically adjusted during training. The activity of the third layer was used to evaluate the model’s pattern separation efficacy.

### Synaptic learning rule

In this work, we introduce a novel synaptic learning rule inspired by early stages of neural development, when neurons receive only a few random synaptic inputs that are insufficient to independently trigger spikes^**60**^. We hypothesize that the synaptic weights of individual neurons are strengthened according to the degree of synchronization among their inputs. In this framework, synchronized synaptic activity directed toward a postsynaptic neuron is assumed to convey meaningful information about the external environment.

Synchronized inputs can encode critical environmental features because their timing strongly influences postsynaptic responses. When multiple presynaptic neurons release neurotransmitters simultaneously, or within a narrow temporal window, the postsynaptic neuron is more likely to fire, thereby providing a reliable mechanism for transmitting stimulus-related information. Such temporal synchronization enables the network to represent more complex signals and supports higher-level processes such as sensory perception, attention, and memory formation. Moreover, neuronal synchrony contributes to oscillatory brain activity, which has been implicated in large-scale functions including motor control, cognition, and decision-making.

To formalize this non-Hebbian learning rule, we first compute the degree of synchronization among synapses converging onto a single postsynaptic neuron. Consider *n* presynaptic neurons, each connected by one synapse. Neurons fire stochastically according to their firing probabilities, producing spike trains represented as binary vectors, where “1” denotes a spike and “0” denotes silence. Spiking activity is evaluated over a 100-bin time window (one trial), after which synaptic weights are updated.

The procedure is as follows:

For each time bin, we compute the mean spike activity of all presynaptic neurons (Eq. 1).

These mean values are transformed into a synchronization score for each time bin (Eq. 2).

At each time bin, the presynaptic spike vector is multiplied by the corresponding synchronization score (Eq. 3).

Finally, the mean of these weighted values is calculated for each presynaptic neuron and used to update its synaptic weight at the end of the trial T (Eq. 4).

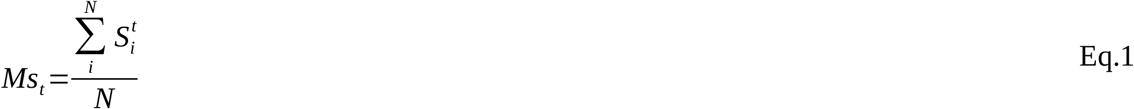

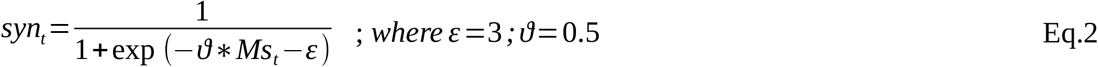

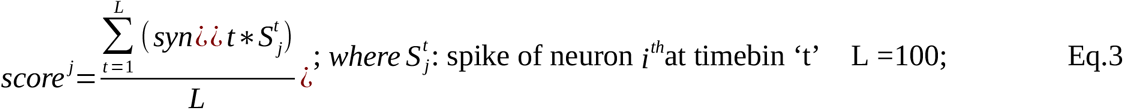

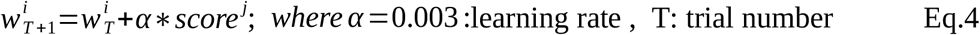

### Neuron model and dynamics

The neuron model used in the proposed feedforward neural network is defined as follows. Neurons in the stimulus layer are predefined probabilistic units, connected to the second layer with an initial connection probability of 0.05. Neurons in the second and third layers are modeled as probabilistic spiking neurons, with their inputs specified by Equation 5.

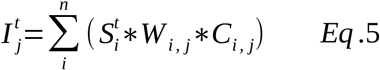

n: number of connected synapses to the single neurons

The probability of spiking for each neuron at a given time bin is determined by Equation 6. Connectivity between neurons and the preceding layer is updated at the end of each training trial according to Equations 7–9. Here, *M*_p_ denotes the average spiking probability of a neuron at the end of a trial.

Equation 8 is inspired by previous work that introduced a probabilistic model of structural plasticity, incorporating the biological relationship between synaptic weight and volume, as well as the dependency of volume on synaptic lifetime^**61**^.

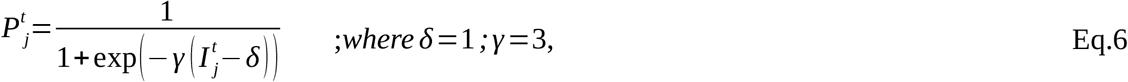

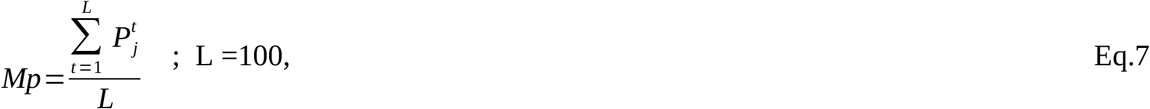

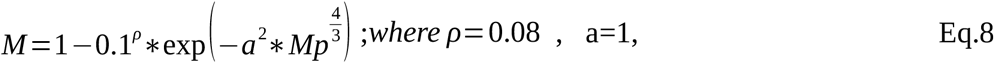

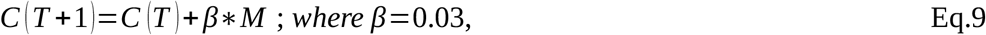

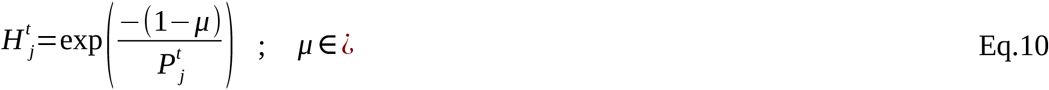

In this framework, *M* represents the probability of forming new synapses and is used to calculate the updated connectivity rate of a single neuron to the preceding layer. As training progresses, modifications to neuronal connectivity alter the total input received by each neuron, which in turn affects its spiking probability, forming a closed-loop mechanism (Fig. 1B).

To stabilize neuronal activity during training, feedback inhibition was implemented. Inhibitory neuron activity is modeled by Equation 10, where 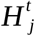 represents the spiking probability of an inhibitory neuron receiving input from a corresponding excitatory neuron at time bin *t*. The parameter *μ* controls the strength of inhibition (Fig. 4A). This inhibition probability is then used to adjust the input activity of the corresponding excitatory neuron.

Thus, both connectivity and spiking probability of each neuron are iteratively updated across trials: connectivity is modified at the end of each trial, while feedback inhibition dynamically regulates excitatory inputs during training.

To evaluate pattern separation efficacy, we adopted the method described in a study on modeling pattern separation in dentate gyrus^**30**^. Specifically, fully separated training input patterns were presented to two independent models. Following training, the evolved connectivity and synaptic weights of excitatory neurons were used to calculate the ordered pattern of average firing rates in the third layer. Pattern separation efficacy under different parameter values was quantified by computing the normalized metric distance between these output patterns.

### Experiments and Simulations

The developed model was evaluated for its pattern separation efficacy under varying levels of inhibitory intensity. We investigated the dynamics of key model parameters, including connectivity and synaptic weights, with particular attention to conditions of network stability and their relationship to separation performance. Furthermore, we examined synchronization within excitatory and inhibitory populations and its contribution to both network stability and pattern separation. These analyses provide important insights into the role of inhibition in cognitive disorders where separation efficacy is impaired, such as autism and schizophrenia.

The proposed PSNN architecture offers a biologically plausible framework for integrating perceptual learning and action selection in autonomous systems. Its event-driven, low-power characteristics make it well-suited for deployment in neuromorphic hardware, with potential applications in cognitive robotics and embodied artificial intelligence. Moreover, the model’s modular and self-organizing nature enables further extensions for complex cognitive tasks such as goal-directed navigation, adaptive behavioral switching, and interaction with dynamic environments.

In addition, the trained Probabilistic Spiking Neural Network (PSNN) was embedded within an artificial agent navigating a two-dimensional simulated environment, employing a random-walk exploration paradigm. The agent’s task was defined as obstacle avoidance, whereby a specific stimulus pattern learned during training was designated as the obstacle to be recognized and avoided. This stimulus was characterized by a unique, non-overlapping activation pattern of neurons, distinguishing it from other benign stimuli with entirely different neural representations.

During navigation, when the agent encountered a stimulus (i.e., came within a predefined spatial proximity to it), the PSNN processed the input and generated an internal representation based on previously learned patterns. If the stimulus matched the learned obstacle pattern, the agent inhibited forward motion and selected an alternative direction, thereby demonstrating recognition and context-dependent decision-making. In contrast, the agent was allowed to traverse areas associated with non-obstacle stimuli, as these elicited distinct neural activity patterns.

Analysis of the third (output) layer of the PSNN revealed distinct population responses to obstacle versus null stimuli, both in terms of mean firing rates and temporal spike distributions (Fig. 10a). The agent performed 1,000 independent random-walk trials, during which its ability to correctly detect and avoid obstacle stimuli was quantified. Stimulus detection was operationalized as the agent’s ability to suppress forward movement upon identifying an obstacle within the spatial threshold (Fig. 10b). Furthermore, detection efficacy was evaluated across varying levels of network inhibition (Fig. 10c). Results indicated that networks trained under moderate inhibitory constraints exhibited the highest obstacle recognition accuracy and navigational performance, underscoring the PSNN’s functional capacity for stimulus discrimination and cognitive control within a dynamic environment. For each parameter setting, we conducted 100 simulations and compared the average results to determine the optimal value.

## Results

To investigate the dynamics of the proposed model and identify optimal conditions for pattern separation, we systematically varied key parameters, placing particular emphasis on the strength of feedback inhibition. All experiments and simulations were performed using MATLAB (MathWorks Inc.), which provided the framework for neural network modeling, data analysis, and visualization. Across all simulations, feedback inhibition played a pivotal role in maintaining network stability by limiting excessive synaptic growth and connectivity during training. It also significantly contributed to enhancing the network’s ability to separate overlapping input patterns.

We found that moderate levels of inhibition consistently provided the best balance between excitatory and inhibitory activity. Specifically, moderate inhibition was most effective in controlling inter-layer connectivity and regulating synaptic plasticity. In contrast, low inhibition led to excessive synaptic strengthening and saturated connectivity, while high inhibition overly suppressed network activity, thereby reducing learning capacity. Networks trained under moderate inhibition exhibited stable and balanced dynamics throughout training (Fig. 2).

Figure 3 depicts the evolution of key network parameters under three levels of inhibitory intensity, focusing on neurons in the third layer; similar trends were observed in the second layer. Connectivity and synaptic weights evolved independently for each neuron in response to presented input patterns.

**Fig. 2.**
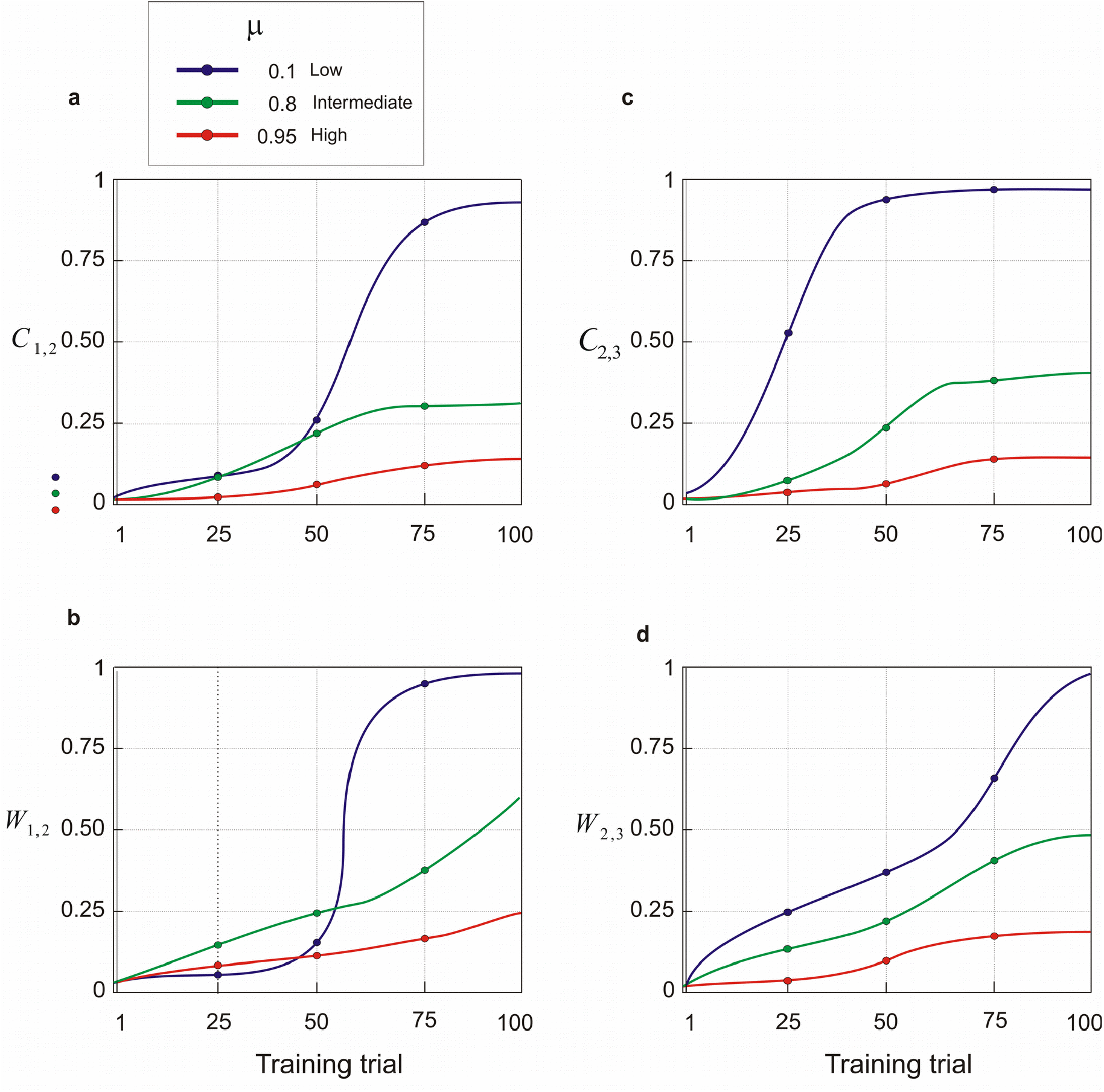
Dynamics of model parameters. Shown are changes in the mean connectivity rate across layers and mean synaptic weights under different levels of feedback inhibition. Simulation results are presented for three inhibition intensities: low (μ = 0.1), intermediate (μ = 0.8), and high (μ = 0.95). Low inhibition leads to uncontrolled growth of both connectivity and synaptic weights across trials, while high inhibition suppresses their increase. Intermediate inhibition achieves balanced dynamics, allowing controlled growth of connectivity and synaptic strength.

**Fig. 3.**
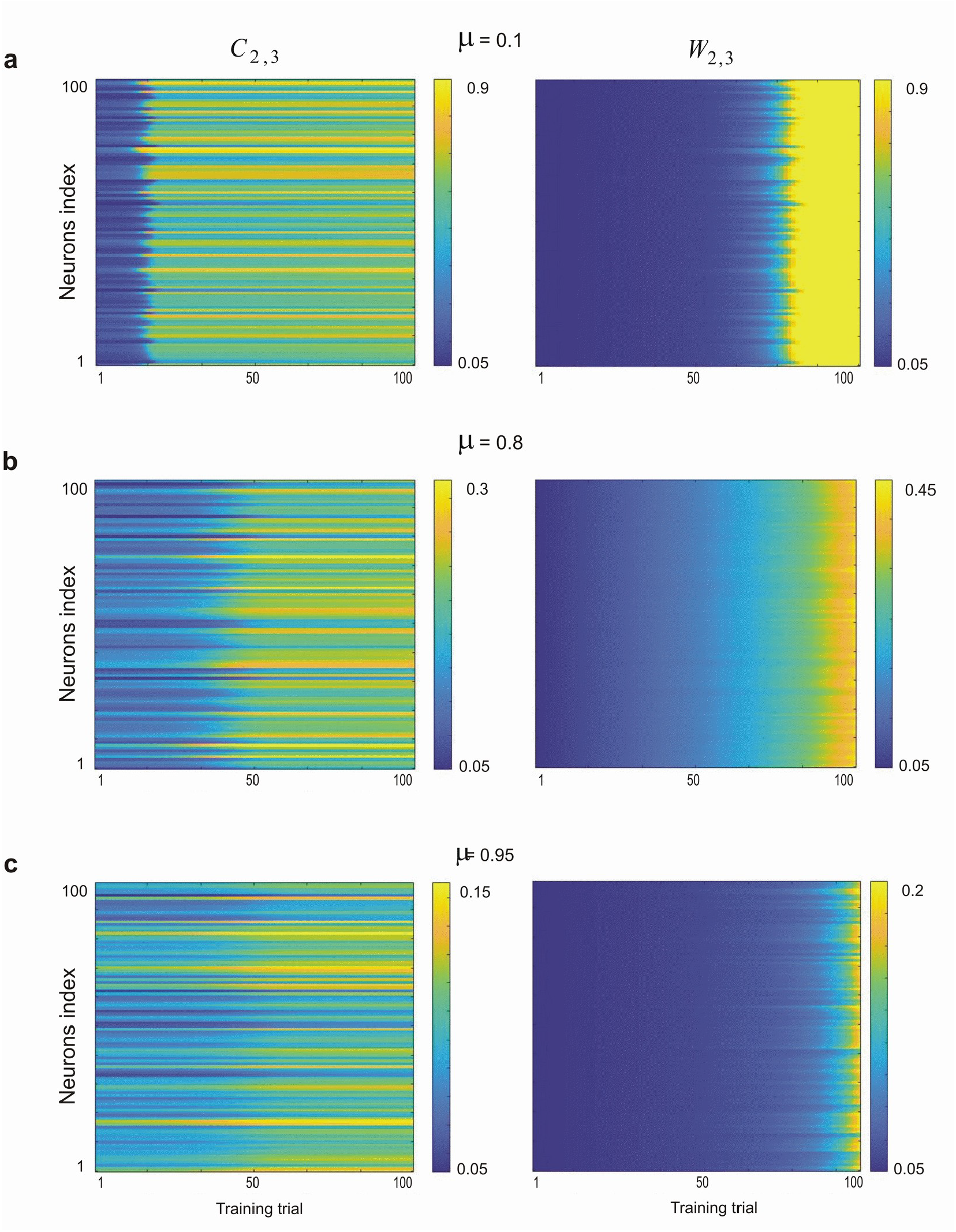
Changes in synaptic weights and connectivity of the third layer across training trials under three levels of inhibition intensity. (**a**) Low inhibition (μ = 0.1), (**b**) Intermediate inhibition (μ = 0.8), and (**c**) High inhibition (μ = 0.95). Synaptic weights and connectivity of individual third-layer neurons evolved independently during training, depending. on their respective input patterns

To quantify the relationship between inhibition strength and separation performance, we analyzed average synaptic weights and connectivity of third-layer neurons across training epochs (Fig. 4). Consistently, moderate inhibition produced higher stability and pattern separation efficacy compared to low (Fig. 4a) and high inhibition regimes (Fig. 4c). Notably, the highest separation performance coincided with network stabilization during training (Fig. 4b), indicating a strong link between dynamic homeostasis and functional performance.

**Fig. 4.**
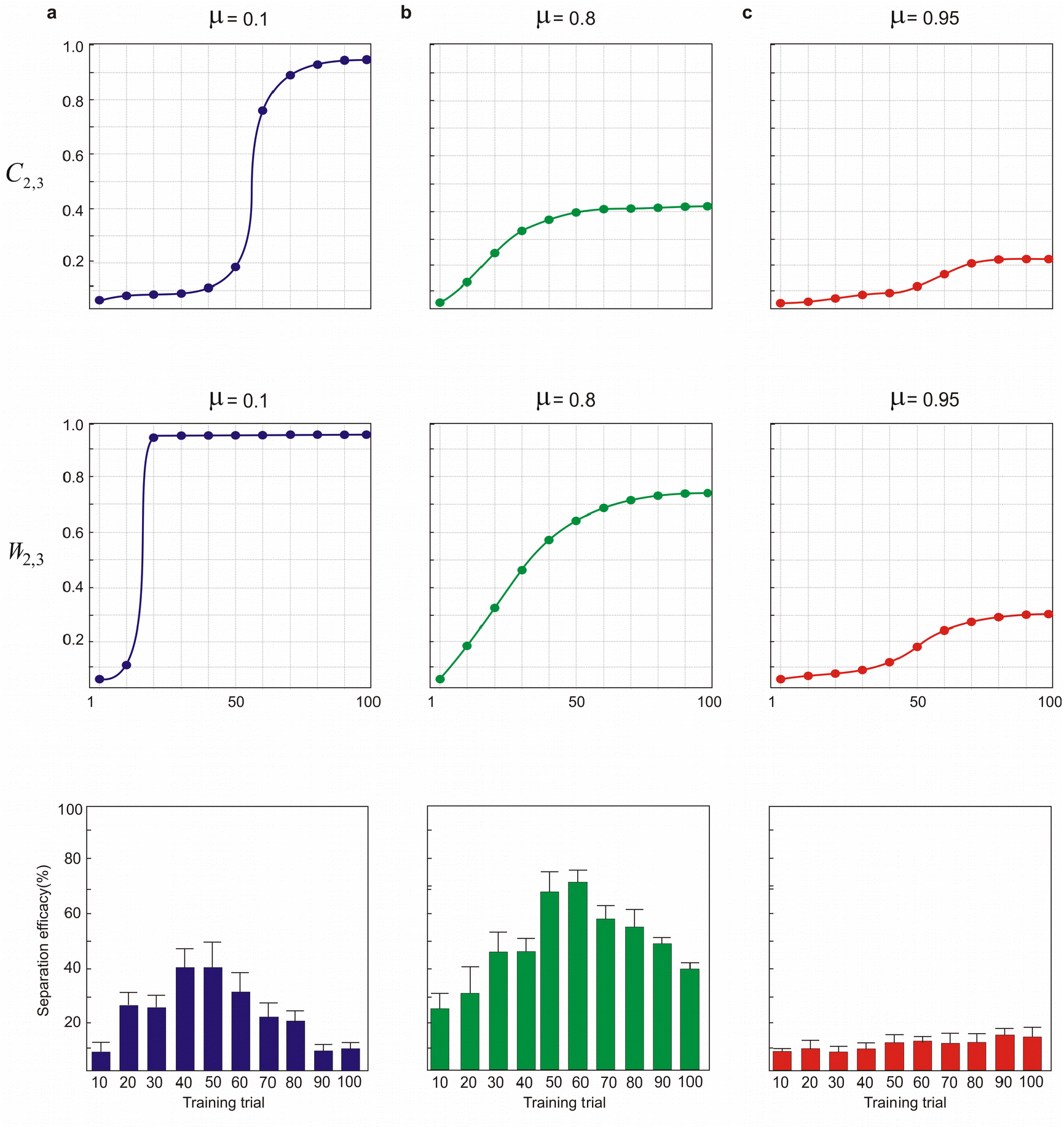
Relationship between pattern separation efficacy and model dynamics across training trials under different inhibition intensities. **(a**) Low inhibition, (**b**) Intermediate inhibition, and (**c**) High inhibition. Results are shown for the average connectivity (C2,3) and synaptic weights (W2,3) of the third layer. The bottom panel illustrates pattern separation efficacy across training trials. Maximum separation efficacy was achieved with intermediate inhibition, around trial 60, when network dynamics stabilized. In contrast, low inhibition produced unstable dynamics characterized by full connectivity and maximal synaptic weights, leading to poor separation. High inhibition excessively suppressed network learning of the input pattern.

Feedback inhibition was modeled via a probabilistic spiking neuron whose firing activity was governed by Equation 9 (Fig. 5a). The inhibitory strength was modulated by the parameter μ, where lower μ values resulted in stronger inhibition and higher values produced weaker inhibition. The spiking rates of neurons in the second and third layers varied systematically with inhibition intensity (Fig. 5b). Importantly, pattern separation efficacy peaked under moderate inhibition conditions (Fig. 5c). Inhibitory neuron activity, as a function of μ, also varied throughout training (Fig. 5d), reflecting the role of inhibition in modulating network dynamics.

**Fig. 5.**
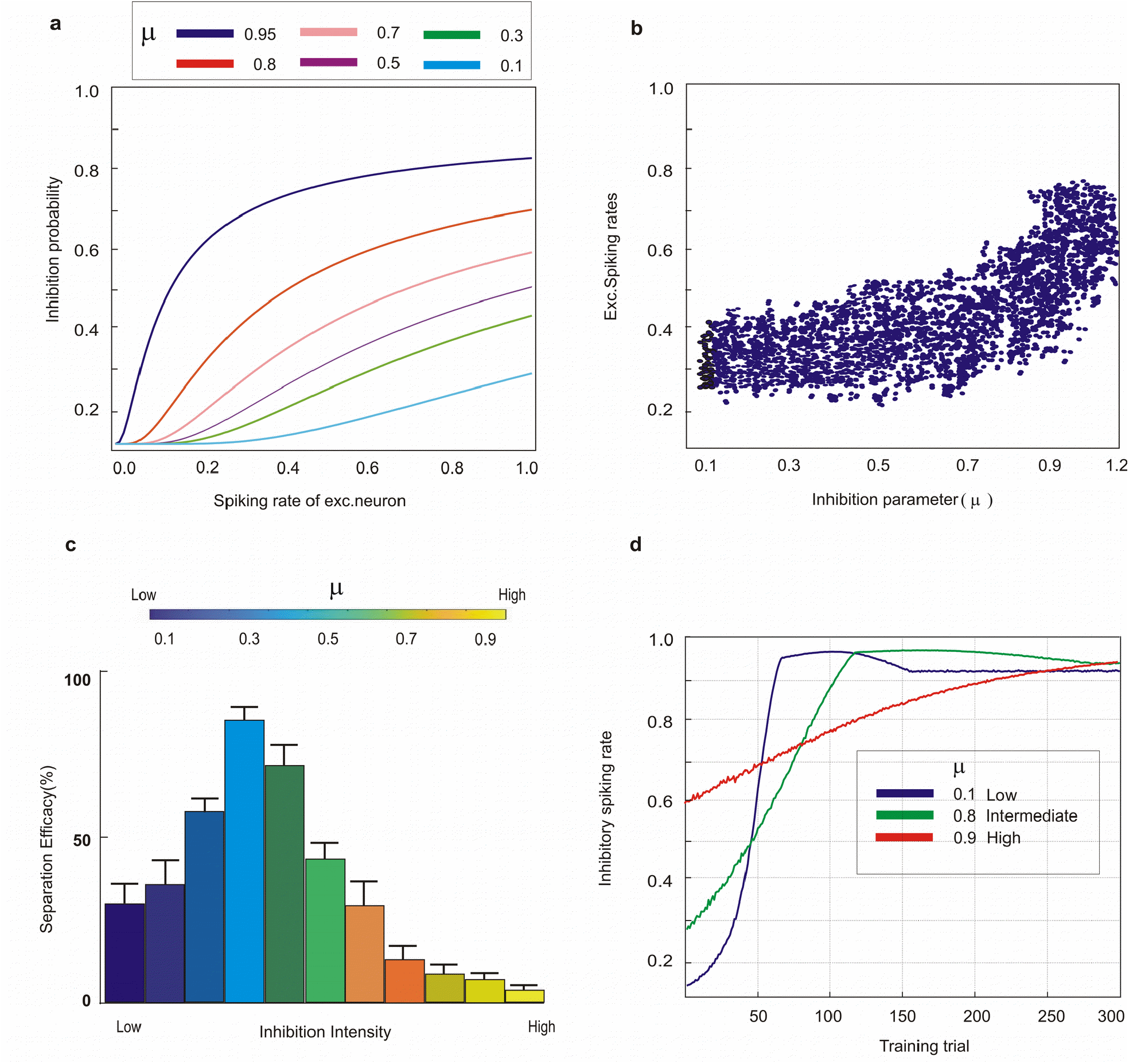
Effects of feedback inhibition on neural activity and pattern separation. (**a**) Feedback inhibition probability at different levels of excitatory spiking activity. Lower α values produce stronger inhibitory activity, while increasing α reduces inhibition intensity. (**b**) Spiking rates of third-layer neurons under different inhibition levels. Weaker inhibition leads to higher excitatory spiking activity. (**c**) Pattern separation efficacy of the model as a function of inhibition intensity. Maximum efficacy is achieved at intermediate inhibition levels.

Figure 6 summarizes the effect of inhibition intensity on layer connectivity and pattern separation. The region highlighted in red corresponds to parameter settings that yielded the highest separation efficacy, reinforcing the importance of maintaining sparse coding, yet functional, connectivity for optimal performance in the developed feedforward SNNs.

**Fig. 6.**
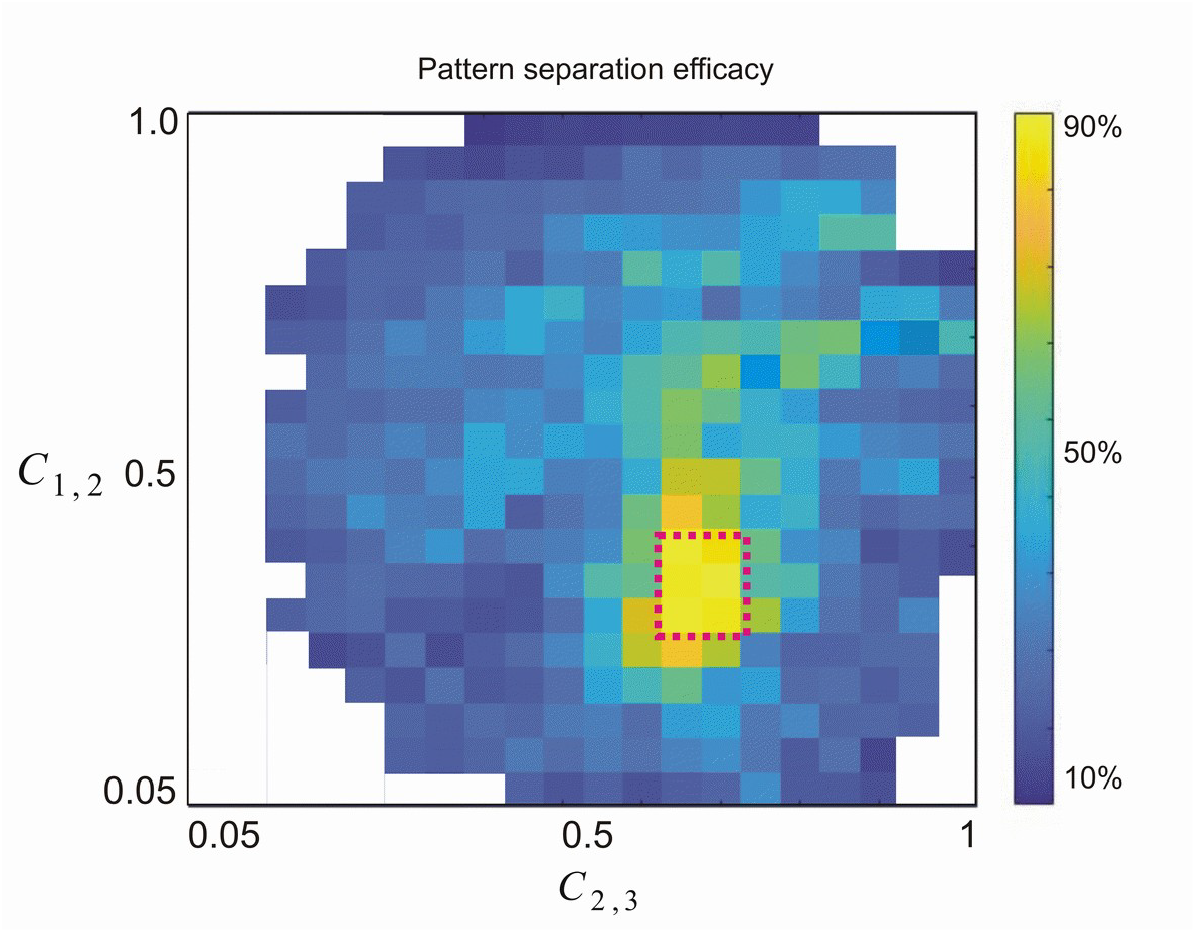
Maximum pattern separation efficacy of the model across different layer connectivity values. Maximum values are highlighted by the dotted red rectangle. Simulations with varying inhibition intensities produced different pairs of connectivity values (*C* _2,3_, *C* _1,2_) at the end of the training phase. These findings underscore the contribution of sparse coding to pattern separation in feedforward spiking neural networks.

To evaluate the contribution of the proposed learning rule, we compared it with a conventional Hebbian plasticity rule. Substituting the synchronization-based learning mechanism with standard Hebbian learning resulted in poor synaptic development during early training, reduced connectivity, and lower separation efficacy (Figs. 7a–d). Across all tested inhibition levels, the synchronization-based rule consistently outperformed the Hebbian counterpart in terms of both network stability and functional separation performance (Fig. 7e).

**Fig. 7.**
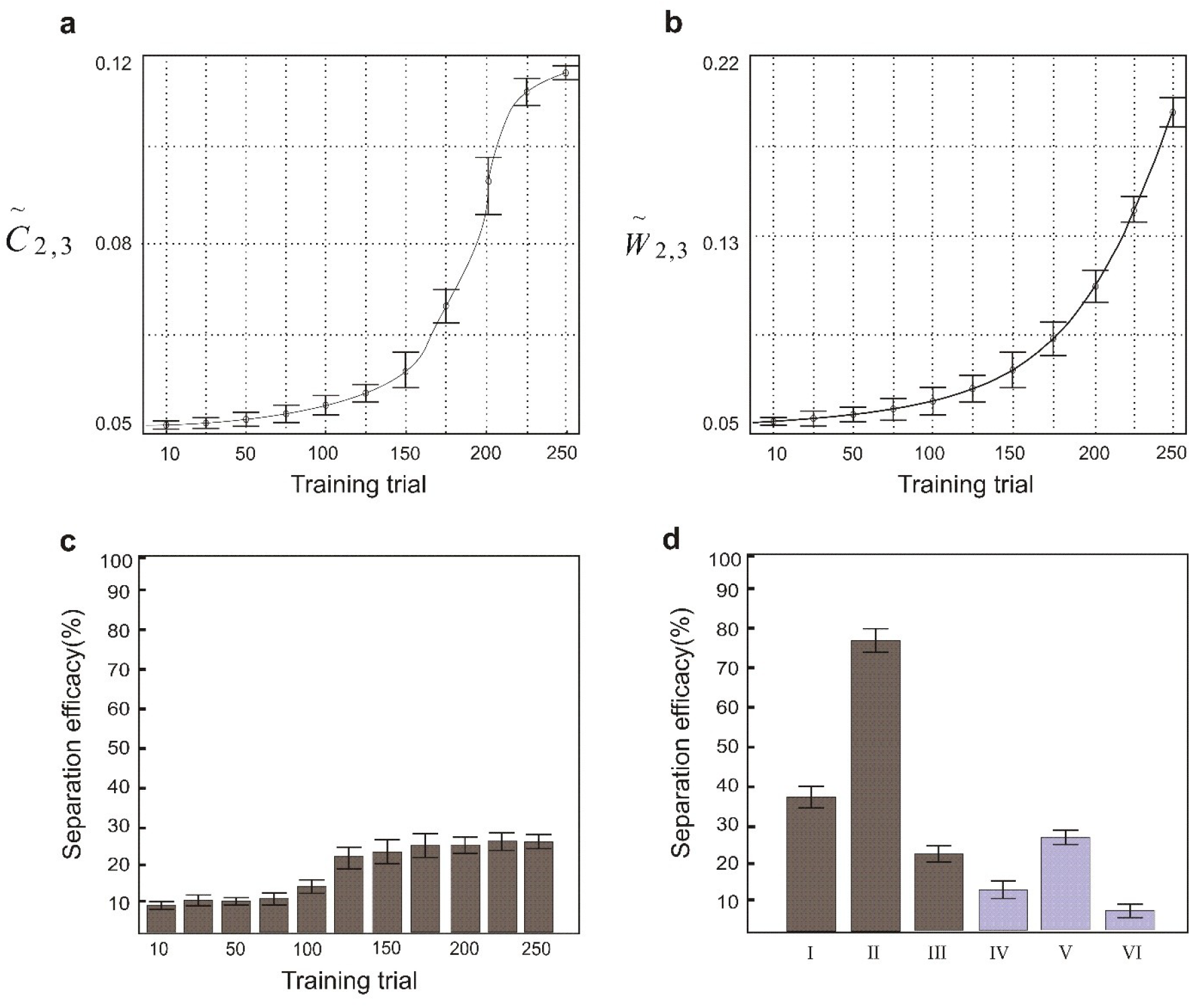
Comparison of the synchronization-based learning rule and the Hebbian learning rule for pattern separation efficacy. (**a**) Connectivity of the third layer across training trials when synaptic weights were updated using the Hebbian rule. (**b**) Mean connectivity rate of the third layer during training, corresponding to panel (**a**). (**c**) Synaptic weights of third-layer neurons across training trials using the Hebbian rule. (**d**) Mean synaptic weight of the third layer, corresponding to panel (**c**). (**e**) Pattern separation efficacy of the model using the Hebbian rule across training trials. All simulations used moderate inhibition intensity (μ = 0.8). (**f**) Maximum pattern separation efficacy of the model under different conditions: I. Low inhibition, synchronization-based rule II. Moderate inhibition, synchronization-based rule III. High inhibition, synchronization-based rule IV. Low inhibition, Hebbian rule V. Moderate inhibition, Hebbian rule VI. High inhibition, Hebbian rule.

We further explored the role of excitatory-inhibitory synchronization in driving separation efficacy. Spiking rates of third-layer excitatory neurons were analyzed under low, moderate, and high inhibition conditions (Figs. 8a–c), with their corresponding average firing rates shown in Fig. 8d. Synchronization dynamics within the excitatory population are presented in Fig. 8e. Notably, peak separation performance occurred when the excitatory and inhibitory populations reached synchronized activity states.

**Fig. 8.**
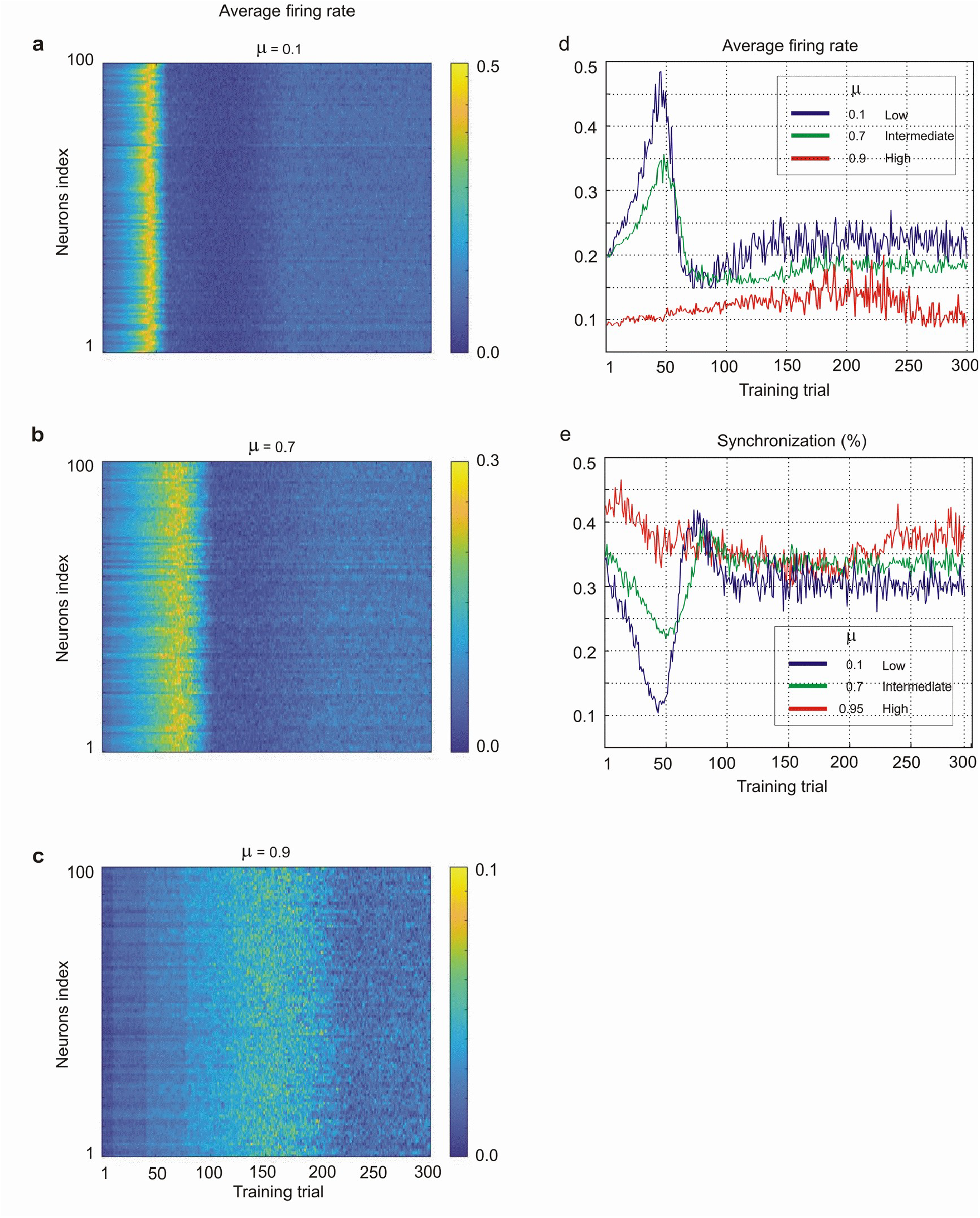
Firing rates and synchronization of third-layer excitatory neurons under different levels of feedback inhibition. (**a–c**) Average firing rates of third-layer excitatory neurons across training trials for (**a**) low inhibition (*μ* = 1.2), (**b**) intermediate inhibition (*μ* = 0.8), and (**c**) high inhibition ( *μ* = 0.1). (**d**) Average firing rate of a representative third-layer neuron across training trials under the three inhibition levels. (**e**) Average synchronization of third-layer excitatory neurons for low, intermediate, and high inhibition. Results show that synchronization increases with training and reaches a maximum, with both the peak value and the corresponding trial number varying by inhibition level. Stronger inhibition leads to lower peak synchronization values reached later in training.

Figure 9a shows the average firing rates of third-layer excitatory neurons across inhibition levels, while Fig. 9b presents the corresponding inhibitory neuron activity. The temporal evolution of excitatory-inhibitory synchrony is depicted in Fig. 9c. Maximum pattern separation coincided with the convergence of excitatory and inhibitory synchrony (Fig. 9d), suggesting that balanced network oscillations are critical for efficient information encoding.

**Fig. 9.**
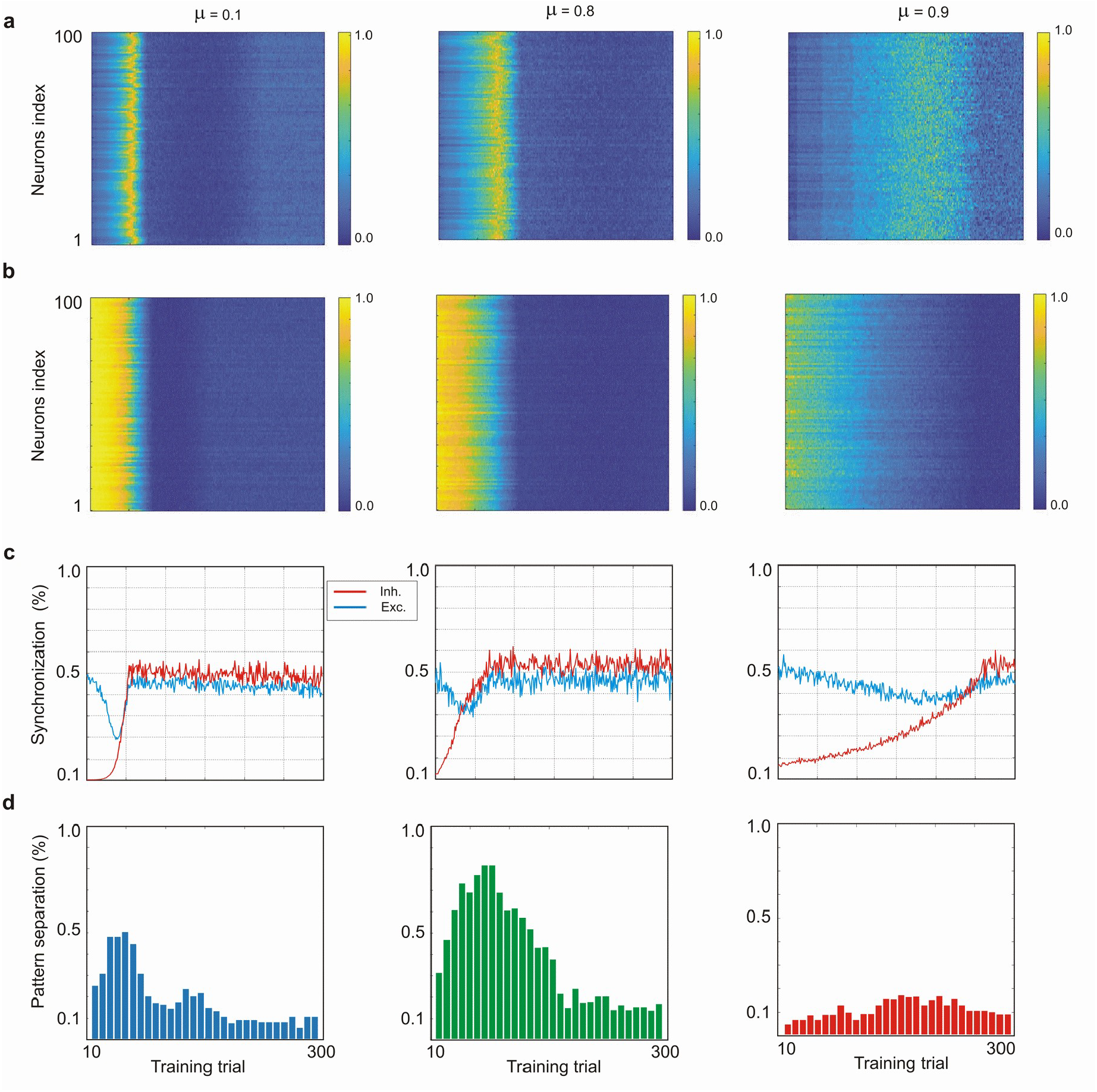
Relationship between pattern separation efficacy and excitatory/inhibitory synchronization. (a) Firing rate dynamics of third-layer excitatory neurons under low, intermediate, and high levels of feedback inhibition. (b) Firing rate dynamics of third-layer inhibitory neurons under the same three inhibition levels. (c) Average synchronization of third-layer excitatory neurons for low, intermediate, and high inhibition. During training, excitatory and inhibitory synchronization gradually converge. Balanced synchronization occurs when excitatory neurons reach their maximum synchronization. (d) Pattern separation efficacy across training trials for different inhibition levels. Maximum separation efficacy is achieved when excitatory and inhibitory populations reach balanced synchronization but declines with further training.

**Fig. 10.**
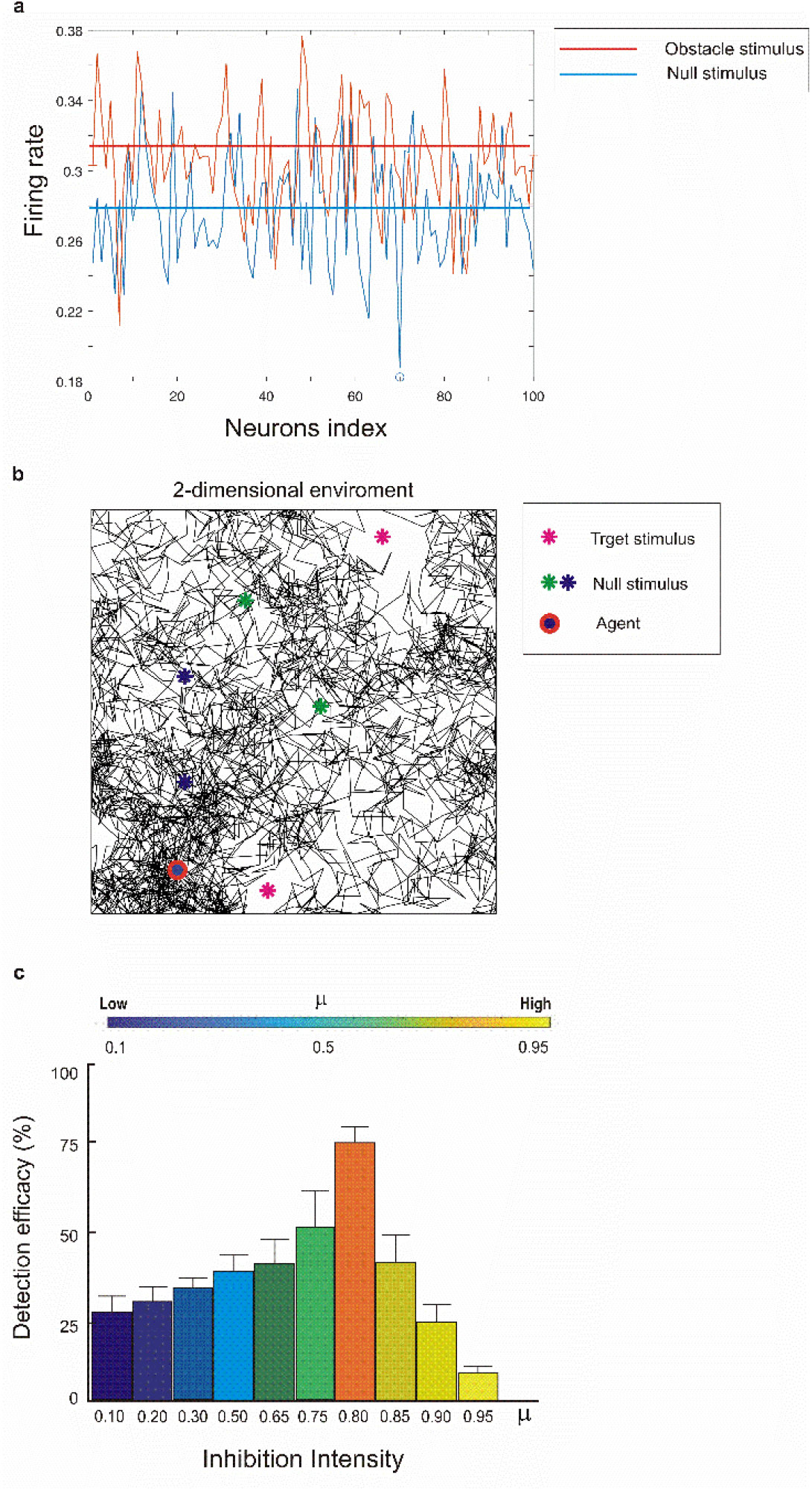
Navigation task using the trained model. (**a**) Firing rates of third-layer neurons in response to the trained stimulus (the obstacle) and an untrained stimulus (null stimulus). Horizontal lines indicate the average firing rates for both cases. (**b**) Navigation of an artificial agent in a two-dimensional random walk (1000 steps shown here). The agent was embedded with the trained model for a specific input pattern as the obstacle. When approaching a stimulus, the agent compared the average third-layer firing rate to a predefined threshold: if below the threshold, the stimulus was classified as null; otherwise, it was classified as obstacle and avoided. (**c**) Detection efficacy of the agent for different levels of inhibition intensity. Maximum efficacy was obtained at intermediate inhibition (μ = 0.8).

These results confirm that the degree of feedback inhibition directly influences network synchrony, which in turn determines separation efficacy.

## Discussion

This study investigates the combined effects of feedback inhibition and a novel synaptic synchronization–based learning rule on self-organization and pattern separation within a probabilistic spiking neural network (PSNN). To the best of our knowledge, this is the first implementation of a synchronization-driven learning mechanism for training artificial spiking neurons. By systematically varying inhibitory strength, we identified a critical regime of moderate inhibition that maximizes separation efficacy, network stability, and dynamic homeostasis. These findings demonstrate that the balance between excitatory and inhibitory populations is essential for optimizing functional connectivity and learning efficiency.

The results indicate that excessive feedback inhibition suppresses a large portion of the excitatory neuron population, ultimately impairing the network’s ability to learn the presented input pattern. Conversely, insufficient inhibition leads to widespread spiking activity among excitatory neurons, which in turn results in dense inter-layer connectivity and the formation of strong synaptic weights. Moderate inhibition produced a well-regulated interplay between synaptic growth and activity, preventing both runaway excitation and excessive suppression. Networks operating under these conditions exhibited sparse but functionally robust connectivity, enabling efficient discrimination of input patterns. In contrast, insufficient inhibition resulted in dense, saturated connectivity and degraded separation, while excessive inhibition hindered learning by silencing excitatory neurons. These results align with neurophysiological observations that the brain maintains excitation/inhibition (E/I) balance to stabilize activity and preserve representational diversity in cortical circuits.

The synchronization analysis revealed that optimal pattern separation coincided with the convergence of excitatory and inhibitory population synchrony. This dynamic equilibrium suggests that precise temporal coordination between neuron types enhances the network’s capacity for information encoding. Such synchronization-driven optimization parallels mechanisms observed in biological systems, where oscillatory coherence supports memory formation, sensory processing, and efficient communication between neural assemblies^**62**^.

Comparative analysis further confirmed that the proposed synchronization-based learning rule outperforms conventional Hebbian plasticity. The Hebbian model exhibited reduced connectivity formation and weaker pattern separation, underscoring the importance of temporal coordination in synaptic modification. By integrating spike-timing information into the learning process, the PSNN captures a more biologically realistic mechanism for adaptive reorganization. This supports the hypothesis that probabilistic and timing-based plasticity rules contribute to the emergence of stable yet flexible network dynamics^**63**^.

The observed relationship between inhibition, synchronization, and separation performance emphasizes the value of viewing learning in spiking systems through the lens of dynamical systems theory. Self-organized balance between excitation and inhibition enables the network to operate near criticality, where small perturbations can efficiently reshape internal representations without destabilizing the overall system^**64**^. This regime facilitates both adaptability and robustness properties crucial for biological cognition and artificial intelligence alike.

From an engineering perspective, these insights provide guidelines for designing efficient neuromorphic and robotic systems. Maintaining moderate inhibitory control and synchronization among functional subpopulations can improve energy efficiency, stability, and online learning capability in hardware implementations. The probabilistic and event-driven nature of the proposed model also favors low-power computation, making it well suited for autonomous cognitive agents operating in dynamic environments.

Taken together, the results highlight that feedback inhibition is not merely a stabilizing factor but a key modulator of learning and representation in self-organizing spiking networks. The interplay between inhibitory control and spike synchronization establishes the structural and functional conditions necessary for efficient pattern separation. Future work will focus on extending this framework to larger-scale cognitive architectures and incorporating reinforcement learning mechanisms to explore adaptive decision-making and continuous learning in biologically inspired robotic systems.

SNNs provide unique opportunities to investigate brain parameters that are not easily accessible experimentally. These include the dynamics of inter-layer connectivity, the biophysical properties of neurons within networks, and the role of synaptic weights in neural computations.

Despite their neurobiological fidelity and event-driven efficiency, SNNs face significant challenges in neuromorphic realization. Local plasticity mechanisms (e.g., STDP) lack efficient gradient-based optimization, constraining large-scale learning and generalization^**65**^.

Neuroscience researchers are especially interested in understanding how neurodevelopment unfolds in biological systems. The mechanisms underlying self-organization such as synaptic plasticity and network optimization for high cognitive performance remain incompletely understood. Cognitive robotics also seeks to mimic neurodevelopment, aiming to build artificial agents capable of exhibiting human-like cognitive skills. Just as infants acquire cognition through gradual experience, modeling these processes may inspire robots that learn adaptively rather than relying solely on preprogrammed skills or supervised algorithms ^**66**^.

In robotics and artificial intelligence, implementing pattern separation in artificial neural networks enhances the ability of robotic systems to recognize and respond to similar but distinct inputs. Pattern separation is crucial for efficiently performing tasks such as object recognition, environment mapping, adaptive decision-making by navigating robots. Algorithms that mimic biological pattern separation allow robots to better generalize from previous experiences while avoiding interference, leading to more reliable learning and behavior in dynamic or ambiguous environments ^**67,68,69,70**^.

Cognitive robotics builds upon these advances by endowing robots with the ability to autonomously perceive, learn, reason, and adapt in complex environments. Central to this approach are self-organizing neural networks computational frameworks modeled after biological circuits and shaped by unsupervised and activity-dependent learning rules^**71**^. Mechanisms such as spike-timing-dependent plasticity (STDP), Hebbian learning, and homeostatic plasticity enable these networks to dynamically self-organize, forming functional representations without external supervision^**72**^. Robotics applications often require real-time decision-making under strict constraints of power, memory, and processing speed. Traditional deep learning models, while powerful, can be computationally expensive and unsuitable for resource-limited robotic platforms, especially mobile or autonomous systems. Developing computationally efficient neural network models addresses this challenge by reducing energy consumption, enabling faster responses, and allowing deployment on lightweight hardware. Such models also support continuous online learning and adaptability, which are critical for robots operating in dynamic and unpredictable environments. By prioritizing efficiency alongside accuracy, robotics can move closer to achieving human-like perception, reasoning, and autonomy in real-world tasks^**73,74,75,76**^.

In practice, self-organized neural networks allow robots to develop internal models that integrate multimodal sensory input, support sensorimotor coordination, and enable higher-level functions such as spatial navigation and decision-making. For instance, spiking self-organizing maps (SOMs) cluster sensory inputs into meaningful feature maps, while recurrent architectures capture temporal dependencies essential for sequential learning. Their adaptability and scalability enhance robotic autonomy, enabling real-time adjustment to variability and uncertainty in real-world tasks through continuous online learning ^**77,78,79**^.

Recent advances combine these networks with neuromorphic hardware, leveraging event-driven computation for real-time processing at low energy cost an essential feature for mobile cognitive robotic systems. Together, self-organized networks and neuromorphic platforms provide a foundation for developing robust, flexible, and cognitively capable robots ^**80**^.

## Acknowledgments

Research supported by ARL Cooperative Agreement W911NF2220143, NIH R01DC019979, NIH R01DC012947, NIH R01NS128924-01, NIH R01MH134118-01

